# Association between the oral microbiome and brain resting state connectivity in schizophrenia

**DOI:** 10.1101/2023.12.22.573165

**Authors:** Dongdong Lin, Zening Fu, Jingyu Liu, Nora Perrone-Bizzozero, Kent E. Hutchison, Juan Bustillo, Yuhui Du, Godfrey Pearlson, Vince D. Calhoun

**Affiliations:** Tri-institutional Center for Translational Research in Neuroimaging and Data Science (TReNDS), Georgia State, Georgia Tech, Emory, Atlanta, GA 30303; Department of neuroscience, University of New Mexico, Albuquerque, NM, 87109; Department of psychology and neuroscience, University of Colorado Boulder, Boulder, CO 80309; Department of psychiatry, University of New Mexico, Albuquerque, NM 87109; Olin Research Center, Institute of Living Hartford, CT 06102, and Departments of Psychiatry and Neuroscience, Yale University School of Medicine, New Haven, CT 06511

**Keywords:** Schizophrenia, Oral microbiome, rsfMRI, functional network connectivity

## Abstract

Recent microbiome-brain axis findings have shown evidence of the modulation of microbiome community as an environmental mediator in brain function and psychiatric illness. This work is focused on the role of the microbiome in understanding a rarely investigated environmental involvement in schizophrenia (SZ), especially in relation to brain circuit dysfunction. We leveraged high throughput microbial 16s rRNA sequencing and functional neuroimaging techniques to enable the delineation of microbiome-brain network links in SZ. N=213 SZ and healthy control (HC) subjects were assessed for the oral microbiome. Among them, 139 subjects were scanned by resting-state functional magnetic resonance imaging (rsfMRI) to derive brain functional connectivity. We found a significant microbiome compositional shift in SZ beta diversity (weighted UniFrac distance, p= 6×10^−3^; Bray-Curtis distance p = 0.021). Fourteen microbial species involving pro-inflammatory and neurotransmitter signaling and H_2_S production, showed significant abundance alterations in SZ. Multivariate analysis revealed one pair of microbial and functional connectivity components showing a significant correlation of 0.46. Thirty five percent of microbial species and 87.8% of brain functional network connectivity from each component also showed significant differences between SZ and HC with strong performance in classifying SZ from HC, with an area under curve (AUC) = 0.84 and 0.87, respectively. The results suggest a potential link between oral microbiome dysbiosis and brain functional connectivity alteration in relation to SZ, possibly through immunological and neurotransmitter signaling pathways and the hypothalamic-pituitary-adrenal axis, supporting for future work in characterizing the role of oral microbiome in mediating effects on SZ brain functional activity.

## 1. Introduction

Schizophrenia, a complex neuropsychiatric disorder, affects around 1% of the world population [1]. It can have enormous influence in personal life and socioeconomics, and can cause large burden to their families and the community. Etiologically, SZ is characterized by a complex interplay between genetics and environmental risk factors. In past decades, collaborative international large scale genetic studies of SZ have enhanced our understanding of genetic risks of the disease [2–4]. However, there is much less effort has been spent on studying the influence of environmental factors or gene-environment interaction in the etiology of SZ. This ‘forgotten’ determinant has been increasingly shown to be relevant in SZ research [5], especially given evidence showing that SZ is a developmental disorder with significant environmental risk factors even during early developmental stages [6, 7]. Thus, studying environmental factors is important to shed light on disease development and pathology.

As an environmental mediator, microbiota are widely colonized in multiple human organs including gut and oral cavity. Colonization can occur at very early life stage (e.g., at birth) and change through the life in response to environmental stimulus such as antibiotic use and diet. Recently, increasing evidence has surprisingly shown “cross-talk” between microbiome and the central nervous system (CNS), that can influence CNS neurodevelopment and function as well as psychological behaviors (e.g., depression, anxiety, cognitive impairment) [8]. There are multiple pathways hypothesized for bidirectional communications between the microbiome and brain including the hypothalamic-pituitary-adrenal axis, neurotransmitters, immune signaling, and recognition of bacterial or host metabolites. Microbial dysbiosis and functional alternations in both gut and oral cavity [9–11], have been demonstrated to involve the regulation of multiple clinical traits, e.g., depression, immune disorders and SZ [12, 13]. Recent landmark findings of gut-microbiome-brain axis have suggested that gut microbiome community can induce inflammation in peripheral and brain system and thus alter behaviors[8]. Certain bacteria (e.g., *proteobacteria*) have been highly suggested to cause the neuro-inflammation via increased brain-blood-barrier permeability and toll-like receptor (TLR4) mediated inflammatory pathways[14–16]. In addition, microbial metabolites may crosstalk with host (epi)genetics and thereby regulate inflammatory process and other metabolic pathways in the process[17, 18]. While the findings suggest the potential role of the microbiome in the pathogenesis of SZ, there are still many unknown regarding the role of the microbiome in SZ, and the relationship of the microbiome to brain function is SZ is unknown.

Characterizing brain functional changes in studying host-microbiome-brain axis will enrich the understanding of the microbial influence on brain function and behaviors. The delineation of the microbiome-brain circuit relationship can help to clarify the relationship of a particular SZ-related microbial alteration on specific brain circuit and function. Neuroimaging is a powerful tool to characterize the structure and function of specific brain circuits and has been applied usefully to brain functional mapping and psychiatric neuroimaging diagnosis [19]. Recently, a few studies have also found the influence of the microbiome on brain circuits related to memory and depression [20–22]. Our recent work has also identified that the abundance of some immune system-related microbiota are significantly changed along with reduced functional connectivity in reward and executive control circuits in smokers [23]. By integrating the oral microbiome with brain activity measures from neuroimaging, we hoped to gain more knowledge of the host-microbiome-brain circuit interactions in SZ.

While the majority of prior studies focused on the link between the gastrointestinal microbiome with brain and other diseases, the oral microbiome has increasingly gained interest as the second largest microbial ecosystem. The oral microbiome also shares microbes with the in gastrointestinal microbiome and is convenient to sample. In addition, some pro-inflammatory microbiota from both oral and gut microbiota show concordant disease associations [24] indicating a potential connection between the two sites in contributing to inflammatory diseases [25, 26]. With these linkages and proximity to the brain, there is the potential for oral dysbiosis, similar to gastrointestinal dysbiosis, to affect brain activity. However, knowledge of how oral microbiome compositional changes might influence neurological signaling in the context of SZ is still limited.

In the current work, we leveraged next generation 16s rRNA sequencing and resting-state fMRI techniques to assess both microbial ecosystem and brain functional activity from SZ and HC individuals to explore the potential associations between oral bacterial populations and neurological signaling (e.g., brain functional connectivity) as well as their relationship with SZ diagnosis to deepen our understanding of those components in the pathology of SZ.

## 2. Material and methods

### 2.1 Participants

All participants were recruited for the Center for Biomedical Research Excellence (COBRE) study and the Glutamate and Outcome in Schizophrenia study [27]. Both were approved by the institutional review board of each site and all participants were provided with written informed consents. There were 213 subjects analyzed in this study, including 110 schizophrenia patients and 103 HCs. Among them, 139 subjects had both microbiome and rsfMRI data as summarized in Table 1. Healthy participants were free of any medical, neurological or psychiatric illnesses and had no current history of substance abuse or dependence. Patients met criteria for DSM-IV-TR schizophrenia, schizoaffective disorder, or schizophreniform disorder by clinical research interviews. No significant difference existed between patients and HCs in age, but there was a marginal difference in sex. Both were controlled as covariates in the subsequent analyses.

**Table 1.** Subject Demographics.

|  | SZ (n=110) | HC (n=103) | P value |
| --- | --- | --- | --- |
| Age(mean±SE) | 37.91±1.34 | 36.1±1.19 | 0.318 |
| Sex(M/F) | 91/19 | 74/29 | 0.071 |
| ***Matched with rsfMRI*** | **SZ (n=59)** | **HC (n=80)** |  |
| Age(mean±SE) | 37.67±1.9 | 38.42±1.33 | 0.741 |
| Sex(M/F) | 52/7 | 61/19 | 0.083 |

Figure 1 shows the analysis workflow for integrating brain functional connectivity by rsfMRI and oral microbiome by 16s sequencing from SZ and HC groups. RsfMRI data were preprocessed by the group independent component analysis (GICA) method (group ICA of fMRI toolbox, GIFT, http://trendscenter.org/software/gift) to derive independent functional networks and calculate functional network connectivity (FNC) matrices [28]. A spatially constrained ICA algorithm using the replicated Neuromark_fMRI_1.0 template as spatial priors was estimated for each subject followed by FNC calculation [29]. Microbiome from saliva tissue were sequenced, followed by a preprocessing pipeline to generate the taxonomic table at different levels (e.g., class and species), to characterize the diversity of microbial community, and to predict functional pathways involved by microbiota. The features (e.g., FNC and taxonomic count) are tested for associations with disease status at difference levels (e.g., taxonomy, pathway) by both univariate and multivariate analysis methods. A comprehensive multivariate method was applied to integrate both datasets to explore the FNC-microbiome correlation and their relation to SZ. In addition, the features selected in multivariate analysis were also evaluated for the classification of SZ patients from HCs by cross-validation.

**Figure 1.**
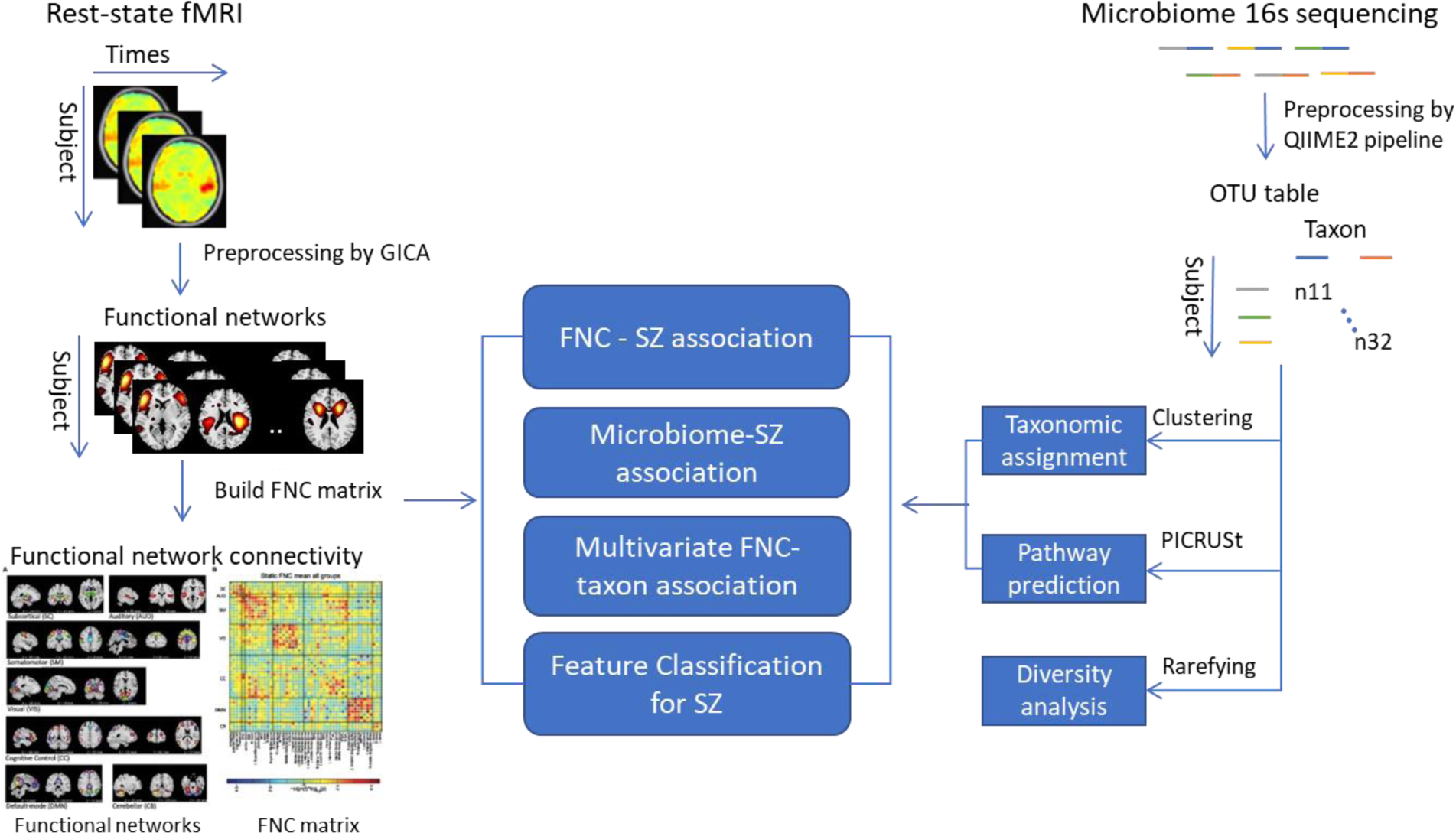
The analysis workflow for resting state fMRI and oral 16s microbiome sequencing data in SZ and HC group.

### 2.2 16S rRNA sequencing

Saliva samples were collected for 16S rRNA amplicon sequencing. Participants provided 5 ml of saliva in a sterile 50 ml conical centrifuge tube and stored in a refrigerator until the DNA was extracted. Sequencing was performed using an Illumina MiSeq covering variable region V4 with primers (5’GGAGGCAGCAGTRRGGAAT-3’ and 5’-CTACCRGGGTATTAAT-3’). Raw sequence data were demultiplexed, followed by quality control by applying the pipeline in DADA2 [30] to generate a number of operational taxonomic unit (OTU) derived by sequences clustering with 97% identity accuracy in mapping to sequences from Greengenes database (http://greengenes.lbl.gov) [31]. Each sequence was trimmed to have a length of 150 base pairs and was aligned by the mafft tool to build a phylogenetic tree [32]. Assigned sequences were further agglomerated at the genus and class levels to have the count table at different taxonomic levels for analyses. All processing scripts were implemented on a QIIME2 (https://qiime2.org/) platform.

### 2.3 Analysis of oral microbiome

Microbiome compositional difference between SZ and HC groups was assessed by comparing cross-sample distance. Raw read counts were first rarefied at 4000 sequences/sample to reduce the sample sequencing bias. Weighted UniFrac distance and Bray-Curtis distance were assessed by the R package ‘vegan’ [33] and tested for group difference by applying permutational MANCOVA (‘Adonis’ function in vegan package) controlling for age, sex. Principal coordinate analysis (PCoA) plots were generated based on the first two principal coordinate vectors (i.e., eigenvectors) from each type of distance matrix to demonstrate the dissimilarity among samples in the 2-D space.

The taxonomic table was normalized to the relative abundances at different taxa levels (e.g., OTU, genus and class). The taxa present in less than 20% of subjects were filtered out, resulting in 82 OTUs, 49 genera, and 16 classes. Each taxon was tested for abundance difference between SZ and HC group by DESeq2 method on raw read count, controlling for covariates age and sex [34]. All of tests were corrected for multiple testing by false discovery rate (FDR, Benjamin-Hochberg method).

A multivariate method, namely sparse partial least square discriminant analysis (sPLS-DA) was applied to evaluate those species (OTUs) contributing to the distinction between the groups [35]. Centred log-ratio transform on relative abundance of OTUs was used prior to multivariate analysis [36]. SPLS-DA method can perform simultaneously variable selection and classification in both data sets (e.g., X and Y data sets) by maximizing the covariance between datasets. The variable selection is achieved by including LASSO ℓ1 penalizations on the loading vector of dataset X, and the classification is by setting dataset Y as the diagnosis vector in sPLS model. Differ to univariate test, sPLS-DA method allows to account for the correlations among the features (i.e, microbiota) during the classification task. The number of features is decided by 10-fold cross validation and 50 repeats of resampling to have the minimum testing balanced error rate. The model with the selected number of features was used for evaluation. Area under the curve (AUC) value is calculated and the weights in the loading vector indicate the contribution of features in the classification.

### 2.4 Analysis of resting state rsfMRI imaging

Participants had resting state functional MRI collected on a 3 T Siemens TIM Trio (Erlangen, Germany) scanner. Images were acquired with an echo-planar imaging (EPI) sequence (TR=2000 ms, TE=29 ms, flip angle=75°) with a 12-channel head coil. Each volume consisted of 33 axial slices (64×64 matrix, 3.75×3.75 mm^2^, 3.5 mm thickness, 1 mm gap). Image preprocessing was performed as previously described [37]. Briefly, this included head motion and slice-timing correction, realignment, co-registration and spatial normalization. The fMRI data were subsequently warped into standard Montreal Neurological Institute (MNI) space using an echo-planar imaging template and then resampled to 3 × 3 × 3 mm^3^ isotropic voxels. The resampled fMRI images were further smoothed using a Gaussian kernel with a full width at half maximum (FWHM) = 6 mm. Afterwards, brain functional networks were estimated by applying the NeuroMark pipeline that leverages a multi-objective optimized ICA with reference (MOO-ICAR) guided with prior network templates [29]. The network templates (neuromark_fMRI_1.0) were derived from a consensus spatial networks by performing ICA on two independent large-sample groups from human connectome project and the genomics superstruct project. 53 components were selected and applied in this study. Time course corresponding to each functional network was filtered using a band-pass filter 0.01–0.15 Hz. Finally, resting state FNC matrix was calculated for each subject based on the correlation coefficients between the time courses of all possible pairs. Each FNC was tested for group difference by multiple linear regression model controlling for covariates (e.g., age and sex).

### 2.5 Linking microbiota with FNC

To explore the relationship between FNCs and taxa, we applied a multivariate method, namely Data Integration Analysis for Biomarker discovery using a Latent cOmponents (DIABLO) [38]. The method is an extended PLS for multiple data sets integration and PLS-discriminant analysis. It is used to explore the correlation of between OTU and FNC through the selection of a subset of features, while showing difference between SZ and HC groups. Similar to PLS-DA analysis, centred log-ratio transform on relative abundance of OTUs was used in multivariate analysis and 10-fold cross validation with 50 repeats of resampling were used for determining the number of features from each data in analysis by using the R package mixOmics v6.6.231. The classification performance by the derived component was also evaluated by computing an AUC value.

### 2.6 Functional pathway analysis of predicted metagenomes

Metagenome content in the samples was inferred from 16S rRNA microbial data, normalized by copy number count to account for the differences of the number of 16S rRNA copies between taxa, and then functional metabolic pathways were predicted based on the MetaCyc Metabolic Pathway Database [39], using Phylogenetic Investigation of Communities by Reconstruction of Unobserved States (PICRUSt) [40]. Pathways were filtered out if they presented on <20% of samples. Next, the group difference of each metabolism pathway between SZ and HCs was tested by Welch’s t-test using Statistical Analysis of Metagenomic Profiles (STAMP) software [41]. Multiple comparisons were FDR-corrected with cut-off set at 0.05 for significance.

## 3. Results

### 3.1 Oral microbiome composition in SZ

Principal coordinate analysis was applied to phylogenetic distance matrices (i.e., weighted UniFrac and Bray-Curtis) to evaluate oral microbiome compositional difference between SZ patients and HCs. As shown in Figure 2, without controlling for covariates (i.e., age and sex), we found significant group difference in weighted UniFrac distance (p= 1.3×10^−3^) and Bray-Curtis distance (p= 0.013) under 9999 times permutation. Differences were still significant (p=1.8×10^−3^ and 0.014, respectively) after controlling for covariates.

**Figure 2.**
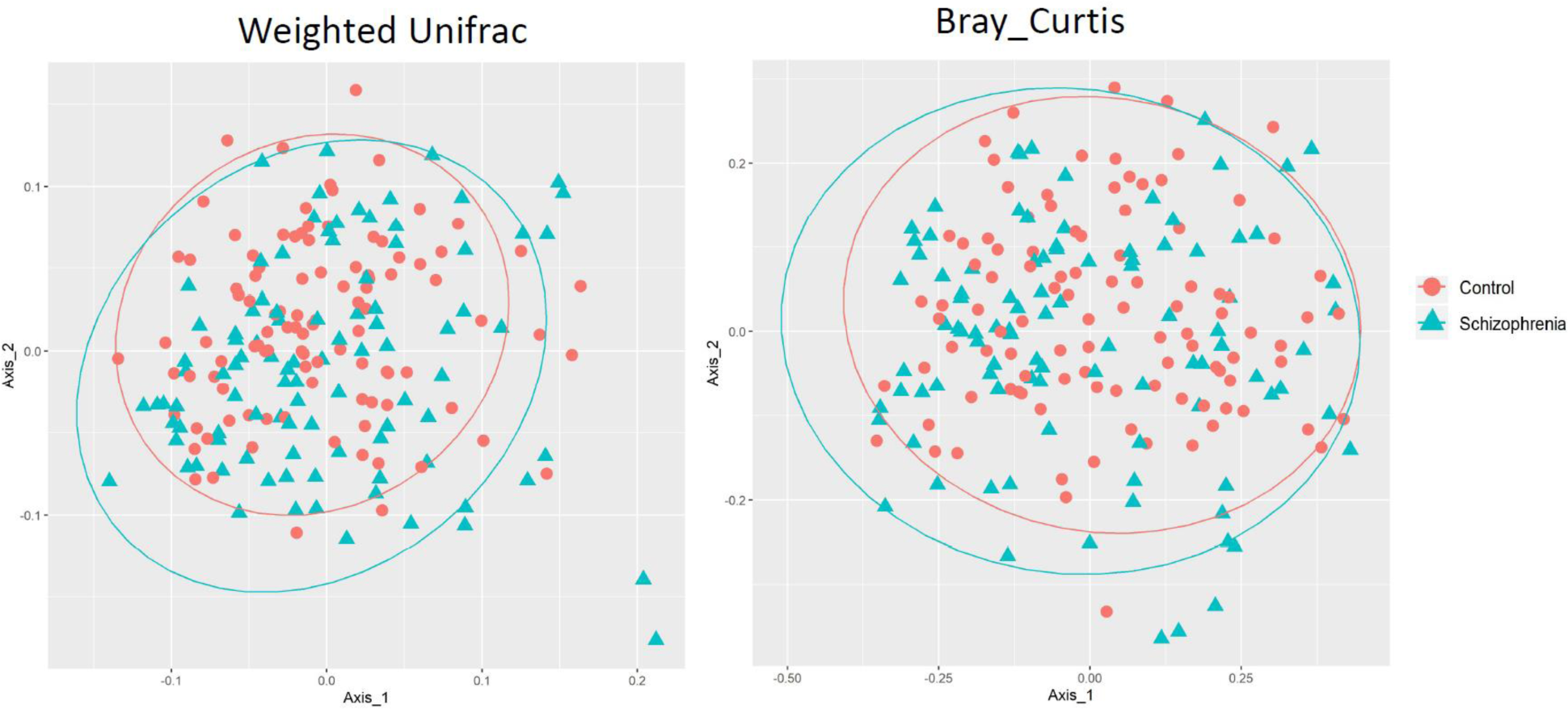
PCoA analysis of microbial composition between SZ and HC groups. Oral microbial composition was evaluated based on (A) Weighted UniFrac distance and (B) Bray-Curtis distance, respectively. Dark blue point indicates the center of eclipse. Axis_1 and Axis_2 are the top two principle coordinate vectors (i.e., eigenvectors) from each distance matrix to show the distances among subjects in 2-D space.

### 3.2 Taxonomic difference between SZ and HC

Group difference of taxa at species, genus, and class levels was also tested by using DESeq2 on read count table. For taxa with significant group difference (FDR≤ 0.05), a multivariate test was applied controlling for covariates (age, sex). Table 2 shows the log-fold change (logFC) and p-value by evaluating the abundance of each taxon between SZ and HC. Significant differences of abundance were identified from 8 classes including 14 species. They mainly consist of Gram-positive bacteria including genera *Actinomyces*, *Rothia*, and *Atopobium* from Actinobacteria, genera *Streptococcus* and *Veillonella* from Firmicutes; and Gram-negative bacteria including genera *Prevotella*, *Porphyromonas* and *Capnocytophaga* from Bacteroidetes, genera *Leptotrichia* from Fusobacteria, and *Lautropia* from Betaproteobacteria. All these bacteria showed higher abundance in SZ compared to HC although with varying logFC: 0.82-1.53.

**Table 2:** List of taxa with significant difference between SZ and HC groups.

| Taxa |  |  |  |  |  | DESeq2 analysis |  |
| --- | --- | --- | --- | --- | --- | --- | --- |
| Phylum | Class | Family | Genus | Species | OTU_ID | logFC | P_adj |
| Actinobacteria | Actinobacteria | Actinomycetaceae | Actinomyces | - | 556682 | 1.86 | $5 \times 10^{-4}$ |
| Actinobacteria | Actinobacteria | Micrococcaceae | Rothia | dentocariosa | 4379034 | 1.21 | $2.3 \times 10^{-2}$ |
| Actinobacteria | Coriobacteriia | Coriobacteriaceae | Atopobium | - | 4451251 | 0.95 | $2.3 \times 10^{-2}$ |
| Bacteroidetes | Bacteroidia | Prevotellaceae | Prevotella | tannerae | 2714267 | 1.53 | $2.5 \times 10^{-3}$ |
| Bacteroidetes | Bacteroidia | Porphyromonadaceae | Porphyromonas | endodontalis | 4303759 | 1.28 | $2.2 \times 10^{-2}$ |
| Bacteroidetes | Bacteroidia | Prevotellaceae | Prevotella | | 4304901 | 0.97 | $2.4 \times 10^{-2}$ |
| Bacteroidetes | Bacteroidia | Prevotellaceae | Prevotella | nigrescens | 4332410 | 1.26 | $1.2 \times 10^{-2}$ |
| Bacteroidetes | Flavobacteriia | Flavobacteriaceae | Capnocytophaga | | 4451646 | 0.94 | $5 \times 10^{-2}$ |
| Firmicutes | Bacilli | Streptococcaceae | Streptococcus | parasanguinis | 4402254 | 0.82 | $2.3 \times 10^{-2}$ |
| Firmicutes | Bacilli | Streptococcaceae | Streptococcus | | 564914 | 0.7 | $5 \times 10^{-2}$ |
| Firmicutes | Clostridia | Clostridiaceae | Peptostreptococcus | stomatitis | 527630 | 1.2 | $2.3 \times 10^{-2}$ |
| Firmicutes | Clostridia | Veillonellaceae | Veillonella | unclassified | 511378 | 1.32 | $2.5 \times 10^{-3}$ |
| Fusobacteria | Fusobacteriia | Leptotrichiaceae | Leptotrichia | | 4430826 | 1.01 | $5 \times 10^{-2}$ |
| Betaproteobacteria | Burkholderiales | Burkholderiaceae | Lautropia | mirabilis | 1084417 | 1.15 | $2.3 \times 10^{-2}$ |

A multivariate method sPLS-DA was further applied to evaluate those species contributing to the distinction between the groups. N=70 OTUs were selected in the final model by 10-fold cross validation. As shown in Figure 3(B), the first component derived from microbiome OTUs can achieve the classification performance AUC = 0.78. Among the top weighted 20 OTUs in loading of component 1, there were 7 OTUs having significant abundance difference between groups, including 5 out of top 10 weighted OTUs from Coriobacteriaceae, Prevotellaceae, Streptococcaceae, Veillonellaceae and Burkholderiaceae.

**Figure 3.**
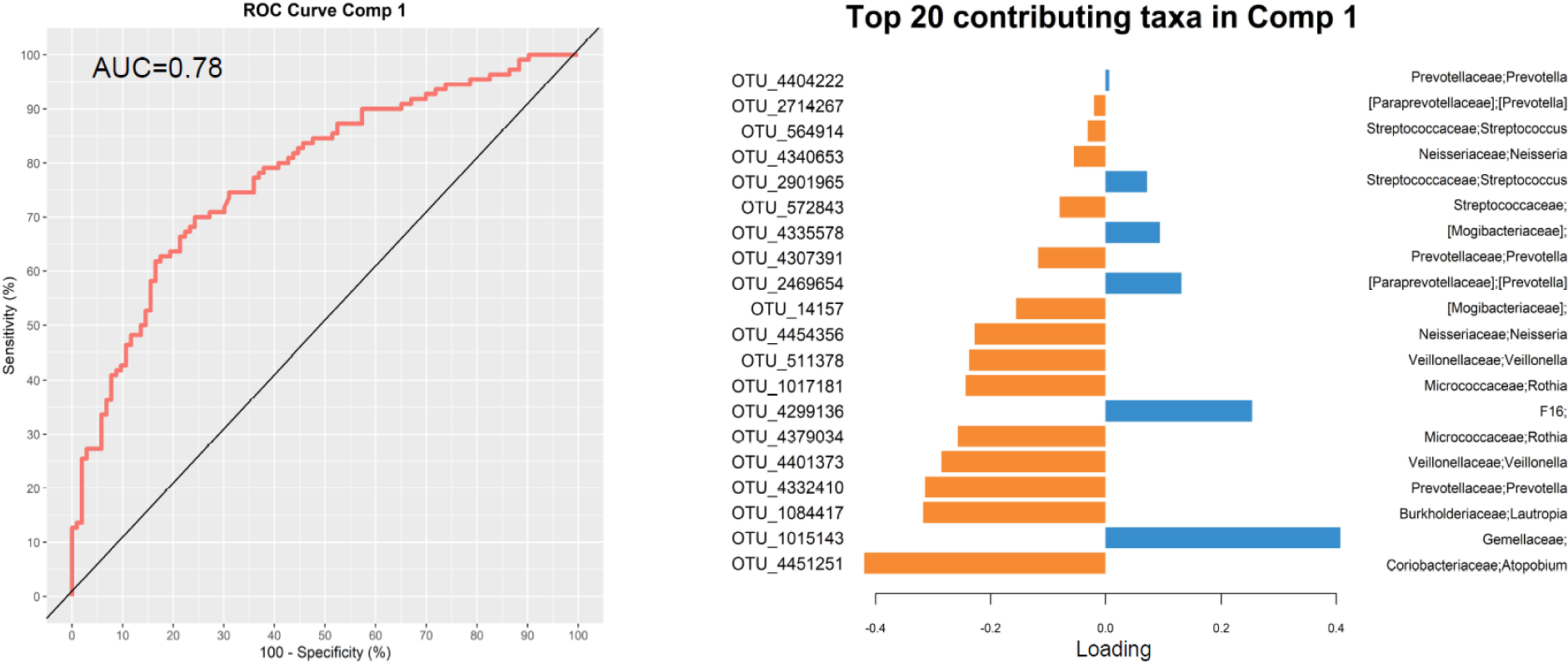
Multivariate analysis on microbiome data at OTU level using sparse partial least square discriminant analysis. (A) plots the ROC curve for classification performance by the first component. (B) lists the weights of top 20 OTUs contributing to the loading from component 1.

### 3.3 Differential FNC between SZ and HC

From multivariate linear regression modeling, we identified 123 FNCs with a significant difference between SZ and HC groups (FDR<0.05) as shown in Figure 4(A). Among these FNCs, reduced connectivity in SZ are mainly involved brain networks from the same domains (e.g., subcortical domain (SC), sensorimotor (SM) domain and visual domain (VIS)), and across domains including SC-cognitive control (CC), SC-cerebellum(CB),and VIS-SM. Increased connectivity in SZ are mainly from cross-domain networks including SC-AUD, SC-SM, SC-VIS and SM-CB. Each brain network derived from GICA analysis was labeled by numeric independent component (IC) index as shown in the triangle boxes of Figure 4 (A). The top brain networks (IC index) that frequently involved those connected functional networks were plotted in Figure 4 (B), including subthalamus/hypothalamus (IC_53) and thalamus (IC_45) from SC domain, superior temporal gyrus (IC_21) from AUD domain, left/right postcentral gyrus (IC_9, IC_11), paracentral lobule (IC_2), and superior parietal lobule (IC_27) from SM domain, middle temporal gyrus (77) from VIS domain, left/right Left inferior parietal lobue (IC_79, IC_81), and precuneus (IC_40) and posterior cingulate cortex (IC_71) from DMN domain.

**Figure 4.**
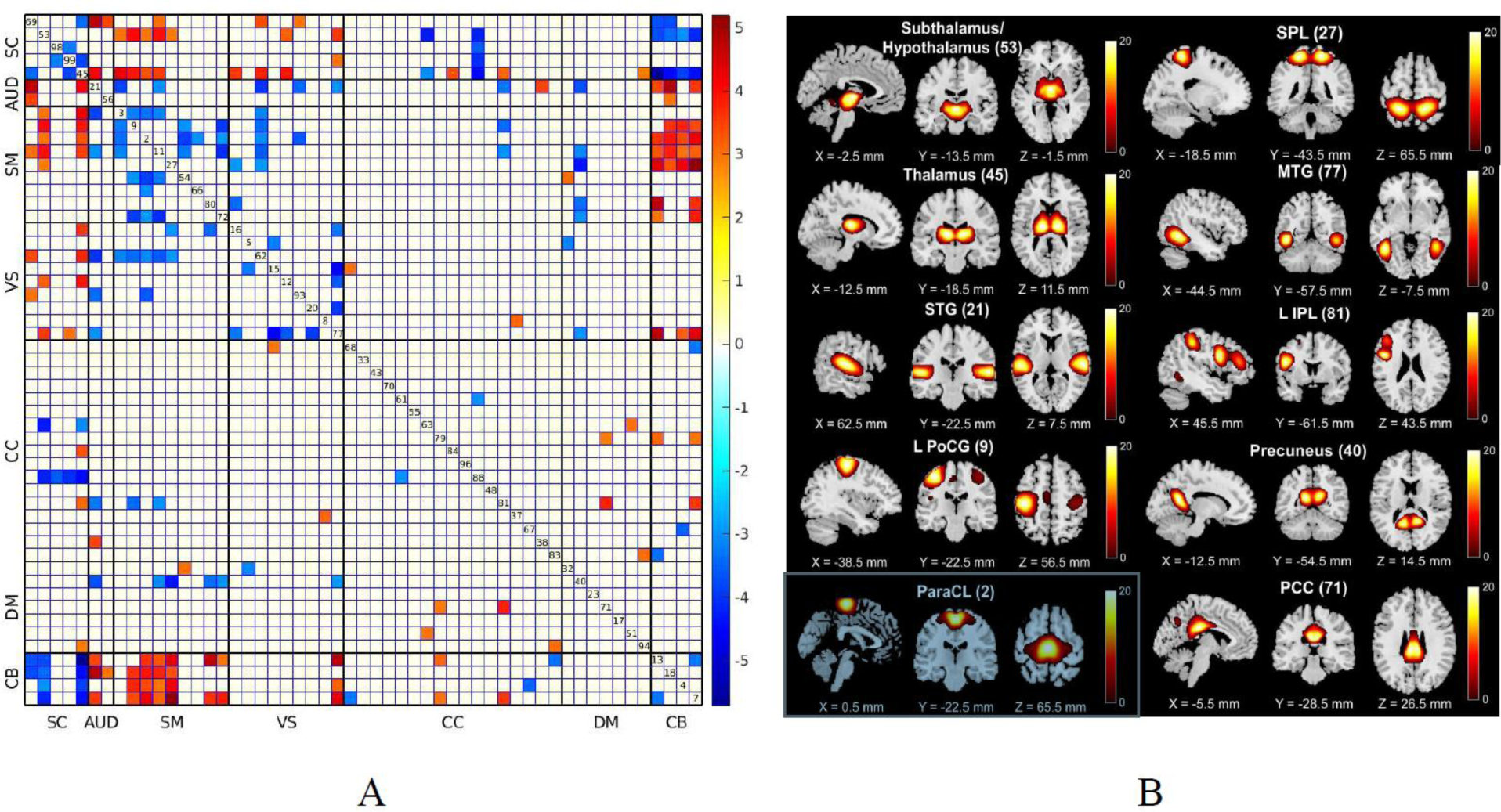
The identified FNCs with significant differences between SZ and HC groups (A) and spatial mapping of top 10 brain networks involved in the connectivity.

### 3.4 Multivariate analysis linking FNC and microbiome

To test for the associations between microbiome abundance and brain functional connectivity strength as well as performance in classifying SZ from HC, we applied the multivariate method DIABLO to the OTUs and FNCs with supervision by diseases status. Ten-fold cross-validation with 50 repeats was used to decide the optimal number of features of each dataset (20 OTUs and 120 FNCs) for association analysis. As shown in Figure 5 (A), a correlation r = 0.46 was achieved for the first pair of components from both data, although the correlation is higher in SZ than HC group. Pair-wise correlation between the selected OTUs and FNCs was tested and those with correlation value r >=0.3 were shown in Figure 5 (B), including 77 links among 4 OTUs and 35 FNCs. There were 3 OTUs (Actinobacteria:Atopobium; Firmicutes:Veillonella; and Proteobacteria:Lautropia) and 35 FNCs with the high correlations (cut-off = 0.3) also showing significant group difference (Table S1). It can be seen that positive correlation was mainly between OTUs and FNCs from SC and task-positive processing brain networks (i.e., SM, VIS and CB) while negative correlations were mainly between OTUs and FNCs from SC and CB brain networks. In Figure 5 (C), there is clearly two clusters from the selected features (OTU and FNC) showing different patterns between SZ and HC groups. Among them, 7 out of 20 OTUs and 108 out of 120 FNCs show significant group difference by univariate tests. In addition, based on the first component from OTU and FNC, we can achieve the classification performance AUC– = 0.835 and 0.871, respectively, as shown in Figure 5 (D).

**Figure 5.**
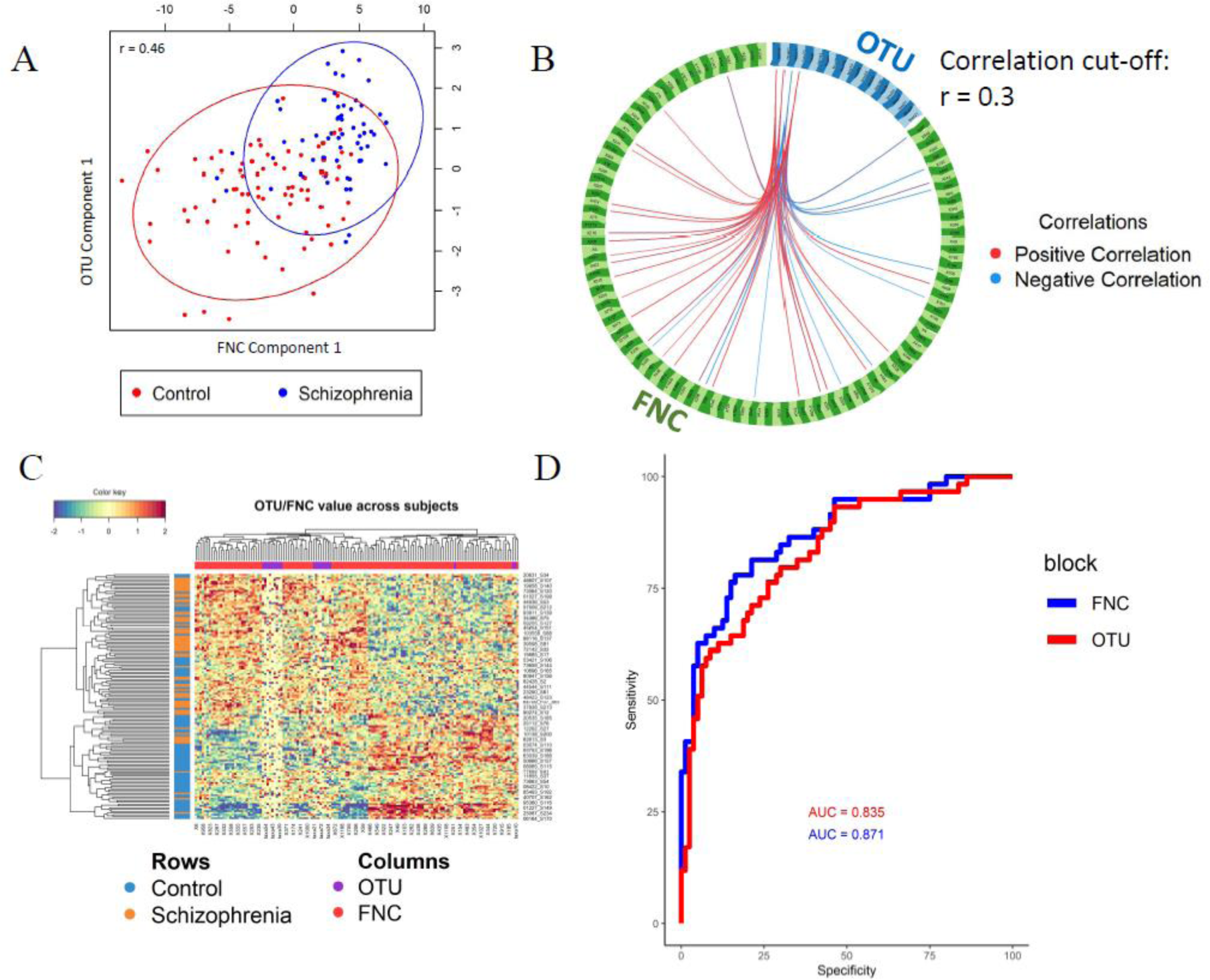
Multivariate method for integrative analysis of both microbiome (OTU) and fMRI (FNC) data. (A) shows the first pair of OUT and FNC components with the correlation r = 0.46 and the distribution of subjects by both components. (B)shows the high correlations (cut-off r = 0.3) between OTUs and FNCs selected from the first pair of components and (C) plots the relative abundance of OTUs and strength of FNC selected in the first pair of components. (D) shows the AUC value of classification by using the first component of OTU and FNC, respectively.

### 3.5 Functional metabolism pathway prediction

There were 384 metabolism pathways predicted at level 3 from MetaCyc database. From these, pathways were removed due to presence in less than 10% of samples. Among the MetaCyc pathways predicted for microbial function, we identified 16 MetaCyc pathways showing significant difference in abundance between SZ and HCs, after correcting for multiple comparisons with FDR thresholded at 0.05, as shown in Figure 6. Depleted pathways were involved with biosynthesis (Kdo2-lipid A biosynthesis and norspermidine biosynthesis), anhydromuropeptides recycling I, L-fucose degradation I, N-acetylglucosamine, N-acetylmannosamine and N-acetylneuraminate acid dissimilation. Enriched pathways were involved with Bifidobacterium shunt, heterolactic fermentation, UDP-N-acetyl-D-glucosamine biosynthesis, mycolyl-arabinogalactan-peptidoglycan complex biosynthesis, chorismate biosynthesis I, and superpathway of aromatic amino acid biosynthesis.

**Figure 6.**
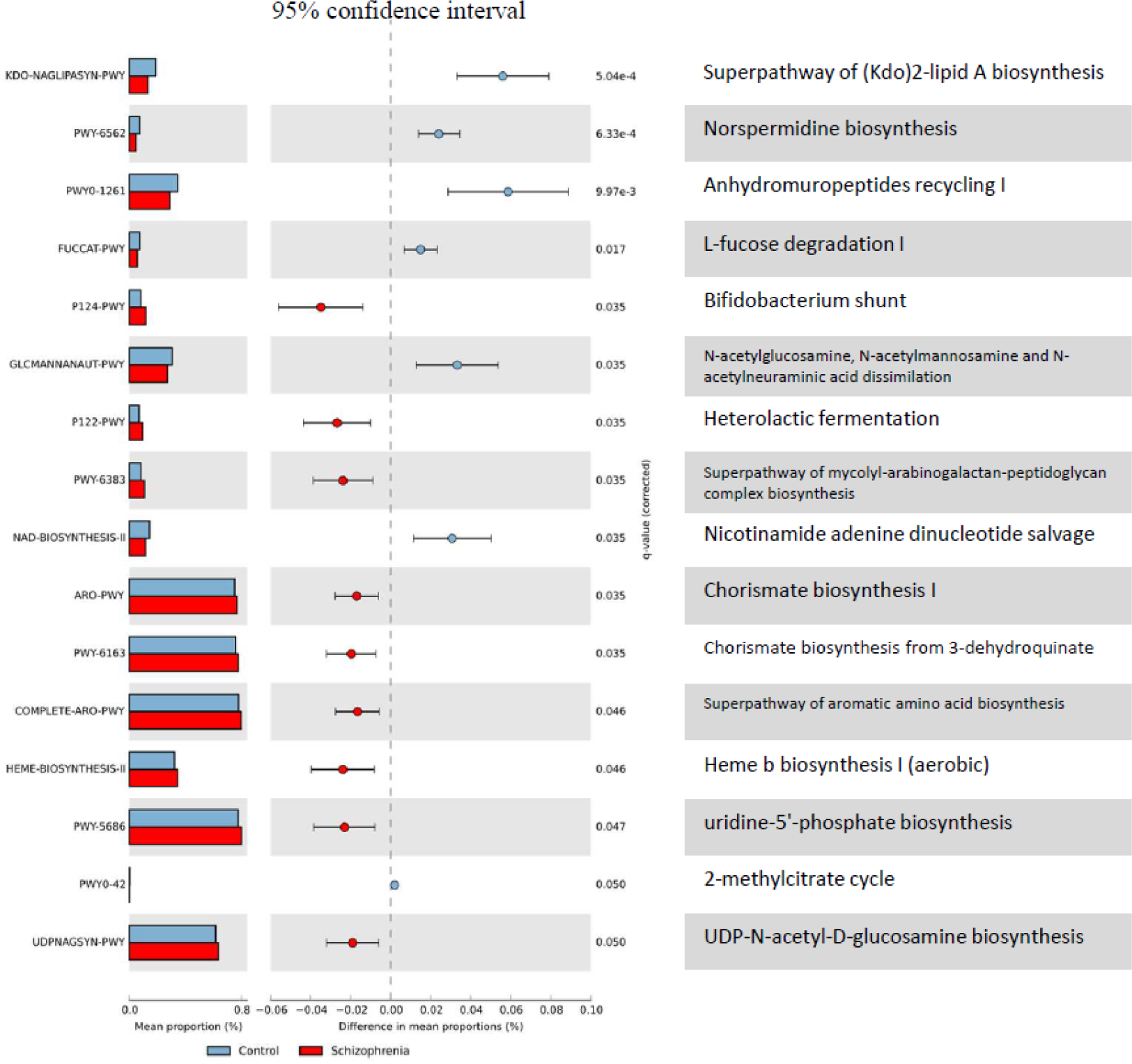
Pathway enrichment analysis based on the predicted metagenomics. The 16 out of 262 MethCyc functional pathways show significant changes in abundance between SZ and HC (FDR < 0.05). Those pathways were predicted from 16S rRNA microbiome sequencing using the PICRUSt algorithm. Mean proportion (colored bar) indicates the relative abundance of the pathway in each group. The difference of mean proportions between groups as well as the 95% confidence interval indicates the effect size of relative abundance change for each pathway.

## 4. Discussion and Conclusions

In this work, we set out to determine whether correlations existed between shifts in the oral microbial population and changes in brain signaling networks in SZ. To achieve this, we used 16s rRNA sequencing to characterize the microbial composition in the saliva of participants (SZ versus HC), and rsfMRI to measure brain functional activity in these same participants. Significant microbial shift was observed in the oral cavity of SZ patients compared to healthy controls, consistent with previously reported findings [42, 43]. In particular, we found the abundance of several microbiota significantly enriched in SZ. Some bacteria were hydrogen sulfide (H_2_S)-producing bacteria including genus *Veillonella*, *Actinomyces*, *Leptotrichia*, *Atopobium*, *Prevotella*, *Porphyromonas* [44, 45]. H_2_S, especially those endogenously produced by gut microbiome have been recognized to play a critical role in human physiology including mediating vasodilation, suppressing oxidative stress in the brain for neuroprotection, and regulating both innate and adaptive immune system [46]. Excess H_2_S and polysulfide production is implicated in the pathophysiology of SZ [47]. Previous studies also shows the enrichment of those bacteria in saliva of schizophrenia patients and suggests that H_2_S-producing bacteria and their product H_2_S are associated with SZ. Although the contributions of those bacteria in schizophrenia is still unknown e.g. they may be trivially associated with secondary disease-related changes in diet, exercise smoking or antipsychotic medication, the findings from present study further suggest a role of H_2_S-producing bacteria in pathology of –schizophrenia. In addition, multiple species form genus *Streptococcus* have been observed in association with inflammatory bowel disease and SZ by modulating pro-inflammatory cytokines secretion among multiple organs [48]. In addition, it is reported that *Streptococcus* species are able to secrete serotonin which is a major neurotransmitter influencing brain activity. A population-based study also suggests that streptococcal throat infections increase the risk of mental disorders, suggesting the host immune intervention [49]. Therefore, significantly increased abundance of multiple species from Streptococcus in saliva of SZ suggests a potential role of this genus in influencing brain function through immunology and neurotransmitter signaling pathways. Genus *Rothia* and *Peptostreptococcus*, as Gram-positive cocci that are widely colonized in the oral cavity, are associated with a wide range of serious infections, especially in immunocompromised hosts [50]. Specifically, studies have shown that species *Rothia dentocariosa* and *Peptostreptococcus stomatis* which are enriched in SZ patients in the present study, could influence the host immune system through stimulating the production of pro-inflammatory cytokines such as TNF-a and IL6 in the oral cavity and gut [51, 52].

Functional enrichment analysis of those predicted pathways of microbiome showed that a variety of associated metabolic pathways were significantly altered in SZ. Several pathways involved in biosynthesis (e.g., Kdo2-lipid A biosynthesis and norspermidine biosynthesis) were depleted in SZ. Kdo2-lipid A serves as the active component of lipopolysaccharide to stimulate potent host immune responses. Significant decrease of Kdo2-lipid A pathway was observed in gut bacteria of SZ in previous study and there was a negative correlation between the pathway with the level of anti-inflammatory cytokine IL10, potentially contribute to a lasting pro-inflammatory state in SZ [53]. Similarly, a nominally significant correlation was observed between Norspermidine synthesis pathway and cognitive dysfunction which is also one of major symptoms in SZ patients [54]. Another set of pathways showing dysfunction in SZ mainly involve glycan metabolism and dissimilation, including anhydromuropeptides recycling I, and degradation of L-fucose, N-acetylglucosamine, N-acetylmannosamine and N-acetylneuraminate which are common carbohydrates making up glycans. Protein glycosylation, which involves the covalent attachment of glycans to proteins, has been reported to be abnormal in SZ or in association with antipsychotic administration [55]. Abnormalities of dysregulated glycosylation have been observed in multiple SZ tissues including blood serum, cerebrospinal fluid, urine, and postmortem brain tissue, potentially representing pathways related to the onset and progression of schizophrenia symptoms [56]. In addition, we also identified pathways upregulated in the oral microbiota in SZ mainly involving glucose fermentation to lactic acid (Bifidobacterium shunt and heterolactic fermentation), biosynthesis of peptidoglycans (UDP-N-acetyl-D-glucosamine biosynthesis and mycolyl-arabinogalactan-peptidoglycan complex biosynthesis) and aromatic amino acids biosynthesis (chorismate biosynthesis I, and superpathway of aromatic amino acid biosynthesis). L-lactic acid is an important metabolite serving as a substance that not only supplies energy to neurons but also can promote neuroprotection through binding to its receptor and suppressing inflammation. as a signaling molecule [57]. Multiple studies have shown d that dysregulation of lactic acid levels is associated with the development of several neuropsychiatric disorders including SZ [58]. It has been suggested that D-lactic acid produced by oral and gut microbiome may diffuse into the blood circulation and further alleviate cognitive and neural functions in patients with SZ [43, 59, 60]. Peptidoglycans have emerged as potential key regulators of gut microbiome-brain interactions, specifically regulating brain development and function by stimulating brain immune responses and modulating neuronal proliferation and differentiation [61]. The aromatic amino acids (e.g., L-phenylalanine, L-tyrosine and L-tryptophan) are the precursors of many neurotransmitters (e.g., dopamine, norepinephrine and epinephrine and serotonin) putatively involved in the pathophysiology of SZ [62]. Collectively, our finding of microbiota enriched in the dysfunctional metabolic pathways related to neurotransmitters production and inflammatory cytokines release indicates the potential path of oral dysbiosis in affecting brain function of SZ.

By correlating microbiome and resting state functional brain imaging data, we found a strong link between a pair of components from each data type. Top microbiota consisting of microbiome component include H_2_S-producing bacteria (*Atopobium* and *Veillonella*) and *Lautropia*. Their increment changes were positively correlated with FNCs mainly between SC (Thalamus, hypothalamus and Caudate), CB and those task-processing networks (SM and AUD), and negatively correlated with FNCs mainly between SC (thalamus and hypothalamus) and CB brain networks. The thalamus and hypothalamus are connected to brain subcortical and cortical regions to form information processing loops. The cerebellum also plays important roles in nonmotor cognitive and affective functions via diverse pathways or interactions with other subcortical or cortical regions such as feed-back and feed-forward loops via the thalamus and pons [46]. A substantial amount of work has observed anatomical and functional deficits of thalamus, hypothalamus, and cerebellar regions in schizophrenia [37, 63, 64]. Our finding of altered connectivity of thalamic and hypothalamus regions with cerebellum and other task processing brain regions in schizophrenia has also been reported in previous studies. In particular, the Cerebellar-Thalamic connectivity has been shown to significantly associated with cognitive deficits in schizophrenia [65]. It is also suggested that bidirectional connections between cerebellar-hypothalamic circuits is important for integrating motor, visceral and behavioral responses such as cognition and emotional expression [66, 67].

The hypothalamic region participates in brain-microbiome interactions through modulation of the hypothalamic–pituitary–adrenal (HPA) axis. HPA axis dysregulation and subclinical inflammation have been reported as a risk factors for conversion to psychosis [68]. Although causal associations are yet to be determined, gut microbiome alterations have been suggested as critical risks for HPA dysregulation through psychosocial stress or by inducing inflammation and blood-brain-barrier permeability impairment [69]. We observed the relationship between the increase of H_2_S-producing bacteria abundance with hypothalamic dysfunction, suggesting an impaired regulation of the HPA axis. Previously, endogenous H_2_S in intestine and brain has been suggested to affect cognitive performance by regulating the release of corticotropin-releasing hormone (CRH) in hypothalamo-hypophyseal portal vasculature to control the secretion of corresponding anterior pituitary hormones and thereby HPA axis [70, 71]. In addition to its effect on the endocrine system, H_2_S seems to serve as an important regulatory mechanism in central nervous system pathologies by regulating brain immune system [46]. A number of studies have recognized the critical role of the inflammatory pathway in microbiome-brain axis where microbial dysbiosis is associated with the release of pro-inflammatory cytokines, altered blood-brain-barrier permeability and neuroinflammation [72]. Our previous study also suggests the association of smoking-induced oral dysbiosis with some brain functional connectivity alternations [23]. Although we did not assess the relationship of H_2_S production between salivary and gut microbiome in this study, our findings suggest the potential role of oral H_2_S-producing bacteria in the brain functional connectivity changes in SZ by HPA axis dysregulation and inflammatory pathways.

The association between the oral microbiome and brain functional network was assessed in SZ. While these associations suggest underlying biological pathways for microbiome-brain link, the causal relationship is not further determined among the features. Given the small sample size and the potential heterogeneity of care and medical treatment received by SZ patients, a replication study with a larger population would strengthen these findings. Additionally, although we tried to control for some covariates in the analysis, other factors such as medications, history of dentition, smoking, alcohol use, and metabolic syndrome (diabetes, obesity, dyslipidemia) could influence the oral microbiome community and brain function and may confound our results. As such, follow on studies should employ stricter criteria for selection of SZ and healthy group are suggested and more clinical metrics need to be evaluated in analysis. Despite these limitations, this study represents the evidence of correlation between population shifts within the oral microbiome and changes in brain function via multiple potential pathways. As only the oral microbiome is accessed in this study, further study comparing the findings in gut microbiome will offer more insights of microbiome-brain interactions in SZ and opportunity for development of novel biomarkers and therapeutic targets for neurological syndromes.

## Declarations

### Ethics approval and consent to participate

Subject recruitment and tissues collection were conducted by the Center for Biomedical Research Excellence (COBRE) study and Glutamate and Outcome in Schizophrenia study at University of New Mexico. The study was approved by the institutional review board of each site and all participants provided written informed consent.

### Additional files

Additional File 1: Supplementary data file listing the correlated (abs(r)>0.3) microbiome and FNC showing group difference between SZ and HC group (FDR<0.05).

